# Sex and estrogen receptor β have modest effects on gene expression in the mouse brain posterior cortex

**DOI:** 10.1101/2020.08.07.241927

**Authors:** Jasmine A Fels, Gabriella A Casalena, Giovanni Manfredi

## Abstract

**Introduction:** Sex differences in brain cortical function affect cognition, behavior, and susceptibility to neural diseases, but the molecular basis of sexual dimorphism in cortical function is still largely unknown. Estrogen and estrogen receptors (ERs), specifically ERβ, the most abundant ER in the cortex, may play a role in determining sex differences in gene expression, which could underlie functional sex differences. However, further investigation is needed to address brain region specificity of the effects of sex and ERβ on gene expression. The goal of this study was to investigate sex differences in gene expression in the mouse posterior cortex, where sex differences in transcription have never been examined, and to determine how genetic ablation of ERβ affects transcription.

**Methods:** In this study, we performed unbiased transcriptomics on RNA from the posterior cortex of adult wild-type and ERβ knockout mice (n=4/sex/genotype). We used unbiased clustering to analyze whole-transcriptome changes between the groups. We also performed differential expression analysis on the data using DESeq2 to identify specific changes in gene expression.

**Results:** We found only 27 significantly differentially expressed genes (DEGs) in wild-type (WT) males vs females, of which 17 were autosomal genes. Interestingly, in ERβKO males vs females all the autosomal DEGs were lost. Gene Ontology analysis of the subset of DEGs with sex differences only in the WT cortex revealed a significant enrichment of genes annotated with the function ‘cation channel activity’. Moreover, within each sex we found only a few DEGs in ERβKO vs WT mice (8 and 5 in males and females, respectively).

**Conclusions:** Overall, our results suggest that in the adult mouse posterior cortex there are surprisingly few sex differences in gene expression, and those that exist are mainly related to cation channel activity. Additionally, they indicate that brain region-specific functional effects of ERβ may be largely post-transcriptional.

## Introduction

Sex differences in numerous aspects of neurobiology and cognitive functions have been described (McEwen, Gray, & Nasca, 2015; Shansky & Woolley, 2016). Sexually dimorphic circuits exist in many brain regions and involve a diverse group of neurotransmitters and neuropeptides (Panzica & Melcangi, 2016). Behaviorally, male and female rodents differ in reproductive behaviors, aggression, social interactions, and learning and memory (Choleris, Galea, Sohrabji, & Frick, 2018). Furthermore, sex differences in the brain are translationally relevant. Women are more susceptible to anxiety disorders, major depression, substance abuse disorders, Alzheimer’s disease, and multiple sclerosis, while men are more likely to develop neurodevelopmental disorders, including autism spectrum disorders, as well as Parkinson’s disease and amyotrophic lateral sclerosis (Zagni, Simoni, & Colombo, 2016).

The molecular basis for sex differences in the brain is still largely unknown, but one candidate is sexual dimorphism in gene expression. In the adult human brain, one study showed over 2,000 genes with differential expression between males and females (Shi, Zhang, & Su, 2016). Conversely, another study with a larger sample size and wider age range found very few differences in quantitative transcript abundance (Trabzuni et al., 2013). In adult rodents, sexual dimorphism in gene expression has been shown in selected brain regions, including the cortex, a brain region heavily involved in nearly all aspects of cognition (Nishida, Yoshioka, & St-Amand, 2005; Xu et al., 2012; Yang et al., 2006). However, sex differences have never been examined in the mouse posterior cortex, which includes the visual, auditory, and somatosensory areas, regions essential for sensation and behavior.

Estrogen and its receptors are likely candidates for a regulatory role in gene expression in the brain. Estrogen readily crosses the blood-brain barrier, or is synthesized by brain aromatase (Ratner, Kumaresan, & Farb, 2019). Estrogen binds two classical estrogen receptors (ERs), ERα and ERβ, as well as a G-protein coupled receptor termed GPER. Evidence suggests that ERβ has notable functions in the brain (Weiser, Foradori, & Handa, 2008). ERβ is a known hormone-responsive transcription factor (C. Zhao, Dahlman-Wright, & Gustafsson, 2010), which regulates transcription directly, by binding to estrogen response elements (EREs) in the enhancer regions of its target genes, or indirectly, by interacting with other transcription factors, most commonly AP-1 (C. Zhao, Gao, et al., 2010). While regulation of transcription by ERβ was previously studied in the female hippocampus (Sarvari et al., 2016) and in the male motor cortex (Varshney et al., 2020), the role of ERβ in regulating sex differences in gene expression in other areas of the cortex has not been investigated.

Several studies have localized ERβ, but not ERα, to neurons and glia in the cortex (Almey et al., 2014; Mitra et al., 2003), including one that used ERβ-GFP transgenic mice to circumvent well-known issues with anti-ERβ antibody specificity (Milner et al., 2010). Furthermore, a large body of evidence suggests that ERβ is important for cortical function. ERβ knockout (ERβKO) mice have impaired social behavior (Choleris et al., 2006), and ERβKO males have deficient motor coordination and electrophysiological abnormalities in the motor cortex (Varshney et al., 2020). Importantly, it was recently shown that central nervous system-specific deletion of ERβ affects social, anxiety, and depressive-like behaviors (Dombret et al., 2020). ERβ also has well-known neuroprotective effects in the cortex, particularly against excitotoxicity and ischemia/reperfusion injury (Azcoitia, Barreto, & Garcia-Segura, 2019; Schreihofer & Ma, 2013; Simpkins, Singh, Brock, & Etgen, 2012). In addition, in the female infralimbic cortex, an ERβ agonist affects synaptic transmission and potentiation (Galvin & Ninan, 2014). Interestingly, ERβ may function differently in the cortex in males and females, as one study showed that treating adult males with an ERβ agonist reduces cortical dendritic spine density (Tan et al., 2012), while another study showed an increase in cortical spine density after treating adult ovariectomized (OVX) females with an ERβ agonist (S. Wang, Zhu, & Xu, 2018). This evidence underscores the importance of studying the effects of ERβ comparatively in males and females, but most studies to date have focused exclusively on one sex. As a result, the contribution of ERβ to sex differences in the brain is largely unknown.

While data suggests that differences in cortical functions exist between males and females, and that ERβ plays an essential role in the cortex, whether gene expression in this region is regulated by sex and ERβ remains to be elucidated. Therefore, in this study we employed global transcriptomic analyses to determine: 1) if sex differences exist in the transcriptome of the adult mouse posterior cortex, and 2) if loss of ERβ affects gene expression in a sex-dependent manner in this brain region.

## Methods

### Animals and Estrus Staging

ERβKO in the C57BL/6J background (Krege et al., 1998) (The Jackson Laboratories stock #004745) and wild type (WT) littermate controls were group housed at a standard temperature with a 12-hour light/dark cycle. Animals were given free access to water and standard chow. Estrous stage was determined with the vaginal cytology method after staining cells with the Fisher HealthCare Hema 3 Manual Staining System, which utilizes the Wright-Giemsa stain, according to (Cora, Kooistra, & Travlos, 2015). All animal protocols were carried out with the approval of the Weill Cornell Medicine Animal Care and Use Committee and were performed according to the Guidelines for the Care and Use of Laboratory Animals from the National Institutes of Health.

### Tissue Collection and RNA Isolation

Mice were sacrificed between 17-19 weeks of age by cervical dislocation after anesthesia and the brain was quickly removed. The right lower quadrant of the cortex, corresponding to portions of the visual, auditory, and somatosensory cortices, was dissected, flash frozen in liquid nitrogen, and stored at −80°C. Cortices were then thawed and homogenized in Trizol, and RNA was extracted with chloroform, then purified using the Promega SV Total RNA Isolation System. Total RNA was analyzed for quality control with an Agilent 2100 Bioanalyzer and quantified with the Thermo-Fisher NanoDrop System, and all samples had RNA Integrity Numbers > 9.

### RNA Preparation and Sequencing

cDNA libraries were prepared from 1µg RNA by the Weill Cornell Genomics Resources Core Facility, using the Illumina TruSeq RNA Sample Preparation kit with oligo-dT beads and uniquely barcoded sequencing adaptors. Libraries of comparable sizes were obtained for all samples and were analyzed for quality using the Agilent 4200 TapeStation and the Thermo-Fisher Qubit Fluorometer. Libraries were multiplexed into a single pool and sequenced with paired-end 50bp reads on the Illumina NovaSeq 6000 in a S1 Flow Cell, at a sequencing depth of 50 million reads. Raw reads were trimmed with cutadapt version 1.18, de-multiplexed with bcl2fastq version 2.19, and aligned to the mouse reference genome GRCm38.p6 with STAR version 2.5.2. Transcriptome reconstruction was done with Cufflinks version 2.1.1. The percentage of aligned reads ranged between 94%-97%, the percentage of uniquely mapped reads ranged between 86%-90%, and the mismatch and indel rates per base for all samples were ≤ 0.1. Transcript abundance was measured in paired fragments per kilobase of exon per million mapped reads (FPKM), generated with Cufflinks.

### Quantitative Real-Time PCR

RNA was extracted as above and quantified with the Thermo-Fisher NanoDrop. 1µg of RNA per sample was used for cDNA synthesis with the Promega Im-Prom II Reverse Transcription System. cDNA was used for quantitative real-time PCR (qPCR) with Thermo-Fisher SYBR Green Master Mix in a Thermo-Fisher Quant Studio 6 Flex. Three transcripts were analyzed, *Scn1a, Lrrc55*, and *Kcnq5*. β-actin (*ACTB*) was used as an internal control for normalization and quantification with the ∆Ct method. Primer sequences used were as follows:

*ACTB forward*: CAAACATCCCCCCAAAGTTCTAC
*ACTB reverse*: TGAGGGACTTCCTCTAACCACT
*Lrrc55 forward*: GTGTCCTTGGAACAGAAGTGAC
*Lrrc55 reverse*: TGAGTCTGGAGGCACAGGTA
*Kcnq5 forward*: CCTACCAAGAAAGAACAAGGGGA
*Kcnq5 reverse*: ATGCGCACTCGCTCCTTAAA
*Scn1a forward*: TAACACTTCAGGGGCTATCGAG
*Scn1a reverse*: AGCACTGTTTGCTCCATCTTG

### Data Analysis

Statistical analyses were done using the DESeq2 package from BioConductor in R, which is based on a negative binomial distribution and uses a normalization procedure based on sequencing depth and biological variance. DESeq2 is advantageous because it offers high sensitivity and statistical power, due to its automatic independent filtering and incorporation of empirical Bayes shrinkage estimations for variance and fold changes (for full details, see (Love, Huber, & Anders, 2014)). We used two models to compare groups, one mono-factorial design separating each sample into one of the four experimental conditions, and one multi-factorial design grouping samples by sex and genotype and incorporating an interaction effect between the two variables. All DEGs shown were found with the mono-factorial model, as no additional DEGs were generated using the multi-factorial model. Genes with less than 10 total raw counts were filtered out before running the DESeq2 model, and all other filtering parameters were kept as DESeq2’s defaults. A Wald test was used to determine statistical significance of differential expression, with the cutoff being a False Discovery Rate of < 5% after Benjamini-Hochberg correction. All data visualization was done in R using the ggplot2, pheatmap, corrplot, and venndiagram packages available from CRAN. Principal component analysis was done with DESeq2, unsupervised agglomerative hierarchical clustering was done using hclust with complete linkage, sample-to-sample distances were calculated with dist, and correlations between FPKM values for individual genes and samples were calculated by cor.test using Pearson’s correlation coefficient. Z scores were calculated from FPKM values for each gene using the standard formula (x-µ)/σ, where x is the sample value, µ is the population mean, and σ is the population standard deviation. Gene Ontology analysis was done with the PANTHER GO Enrichment Analysis tool from Gene Ontology Resource and the gprofiler2 package developed by Raudvere et. al. (Raudvere et al., 2019). Cell type deconvolution was done with the MuSiC package developed by Wang et. al. (X. Wang, Park, Susztak, Zhang, & Li, 2019).

## Results

### Quality control and clustering of our transcriptomics dataset

We performed RNA sequencing on the posterior region of the mouse cortex, including portions of the visual, auditory, and somatosensory cortices (Vanni, Chan, Balbi, Silasi, & Murphy, 2017), from male and female C57BL/6J ERβKO adult mice (17-19 weeks of age) and their WT littermate controls (n = 4 independent biological replicates for each of the four experimental groups). We chose to perform this study in randomly cycling females in order to not bias our results for one particular estrous stage. At the time of sacrifice, two WT females and three ERβKO females were in estrus, one WT female was in diestrus, one WT female was undeterminable, and one ERβKO female was in metestrus.

We found high and largely consistent percentages of uniquely mapped reads across samples, ranging between 86%-90%, and identified 48,303 total genes, of which 22,055 were protein coding genes (Supplementary Data). Our identified transcriptome thus covers 98% of the annotated protein coding genes in the mouse reference genome. We first normalized our count data with a variance stabilizing transformation (VST) and saw low dispersion across genes (Fig 1A) and no obvious outliers among samples (Fig 1B), validating the quality of our sequencing data. Unsupervised hierarchical clustering of sample-to-sample distances revealed no grouping into distinct clusters, indicating that overall variability in gene expression is not more different between groups than within each group (Fig 1C). We next visualized the correlation between the average expression of each gene in each group, against that gene’s expression in other groups. The biologically relevant comparisons are WT female vs WT male, ERβKO female vs ERβKO male, WT female vs ERβKO female, and WT male vs ERβKO male (Fig 1D). Surprisingly, the average expression of each gene was remarkably similar in all comparisons, and the high correlation between the average expression of each gene across each comparison indicates that few genes were differentially regulated. Finally, to confirm that our RNA pool was enriched for cortical RNA, we analyzed the expression of known marker genes for different brain regions. Five marker genes each for cortex, hippocampus, and midbrain were chosen using the differential expression tool from the Allen Brain Atlas In Situ Hybridization Data, by selecting the top five genes listed for each region without appreciable expression anywhere else in the brain. This analysis confirmed that our RNA pool was highly enriched for cortical RNA (Fig 1E). Taken together, these data confirmed the quality of our dataset, while also suggesting that differences in cortical gene expression between males and females, and ERβKO and WT mice, are modest.

**Figure 1.**
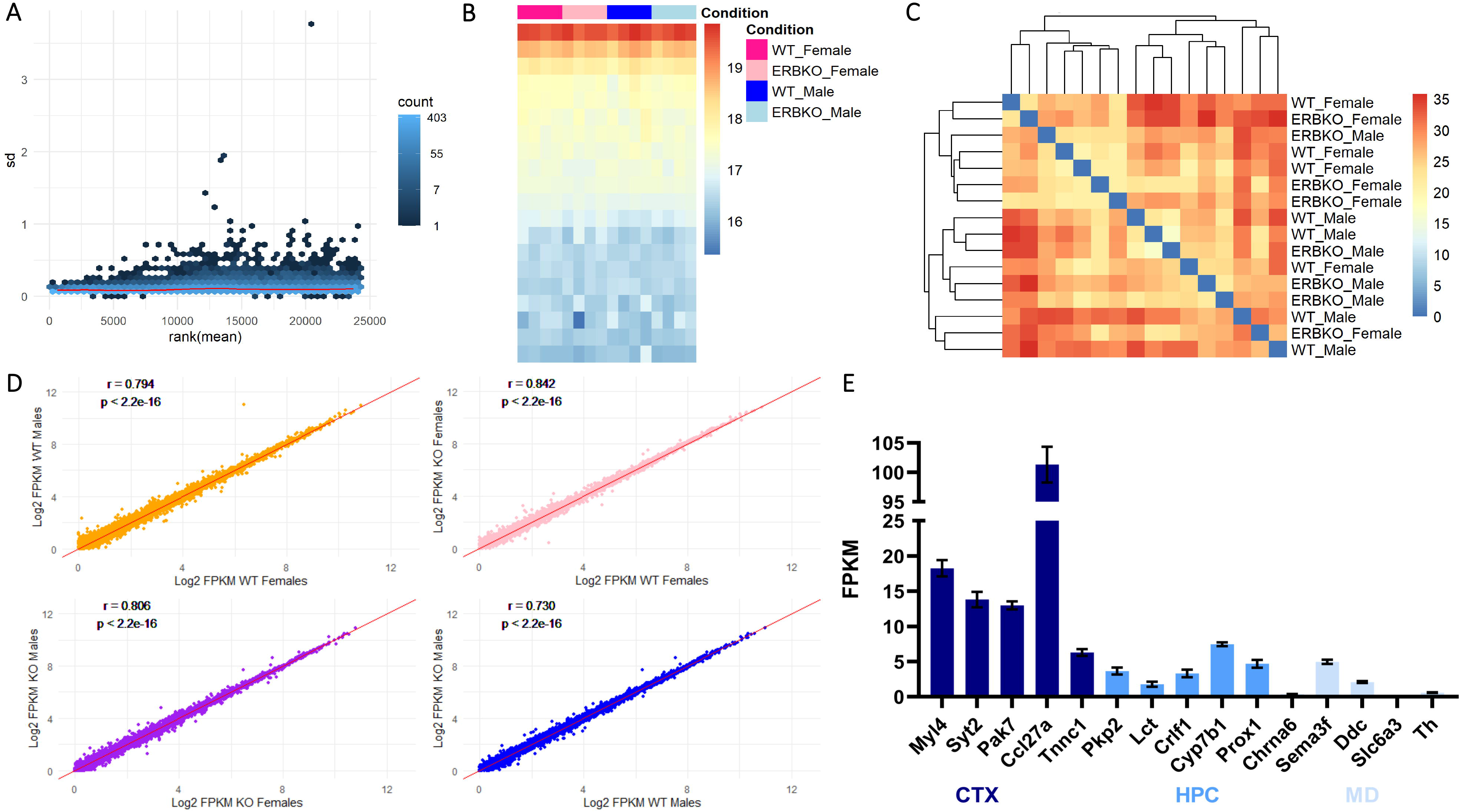
Quality Control of the transcriptomics dataset. **A.** Scatter plot of means ranked in order of magnitude against their standard deviations from variance-stabilized counts. **B.** Heatmap of variance-stabilized counts. Each column corresponds to one sample, and rows correspond to groups of genes ordered by Z-score. **C.** Heatmap of sample-to-sample distances, clustered with unsupervised hierarchical clustering. **D.** Scatter plots of log_2_-transformed FPKM values for each gene with Pearson’s correlation coefficient (r) and p-values shown. The WT female vs WT male comparison is shown in yellow, ERβKO female vs ERβKO male is shown in purple, WT female vs ERβKO female is shown in pink, and WT male vs ERβKO male is shown in blue. This color-coding is continued throughout. **E.** FPKM values showing the expression in our cortical RNA pool of five marker genes each for cortex, hippocampus, and midbrain, averaged for all samples, n = 16. Marker genes were chosen from the Allen Brain Atlas In Situ Hybridization database. Results are presented as mean ± SEM.

One explanation for a lack of transcriptional changes in ERβKO is compensation by another estrogen-responsive receptor. Transcription at EREs is mediated by ERα and ERβ, and while GPER does not directly bind DNA it can participate in estrogen signaling cascades leading indirectly to transcriptional changes (Barton et al., 2018). We therefore looked at the expression levels of ERα, ERβ, and GPER in each group, and found no changes in the expression of ERα or GPER in response to loss of ERβ (Fig 2A), suggesting that there is no compensatory upregulation of either receptor in the ERβKO cortex. Additionally, there were no statistically significant differences in the expression of any of the estrogen receptors between males and females. We also found that expression of ERβ is 11-fold higher than ERα and 6-fold higher than GPER, confirming previous studies showing that ERβ is the most abundantly expressed estrogen receptor in the cortex (Mitra et al., 2003). Furthermore, we found no difference in the expression of the estrogen-synthesizing enzyme aromatase (*CYP19a1*) among any of the groups analyzed, and expression levels of this gene were very low (Fig 2B).

**Figure 2.**
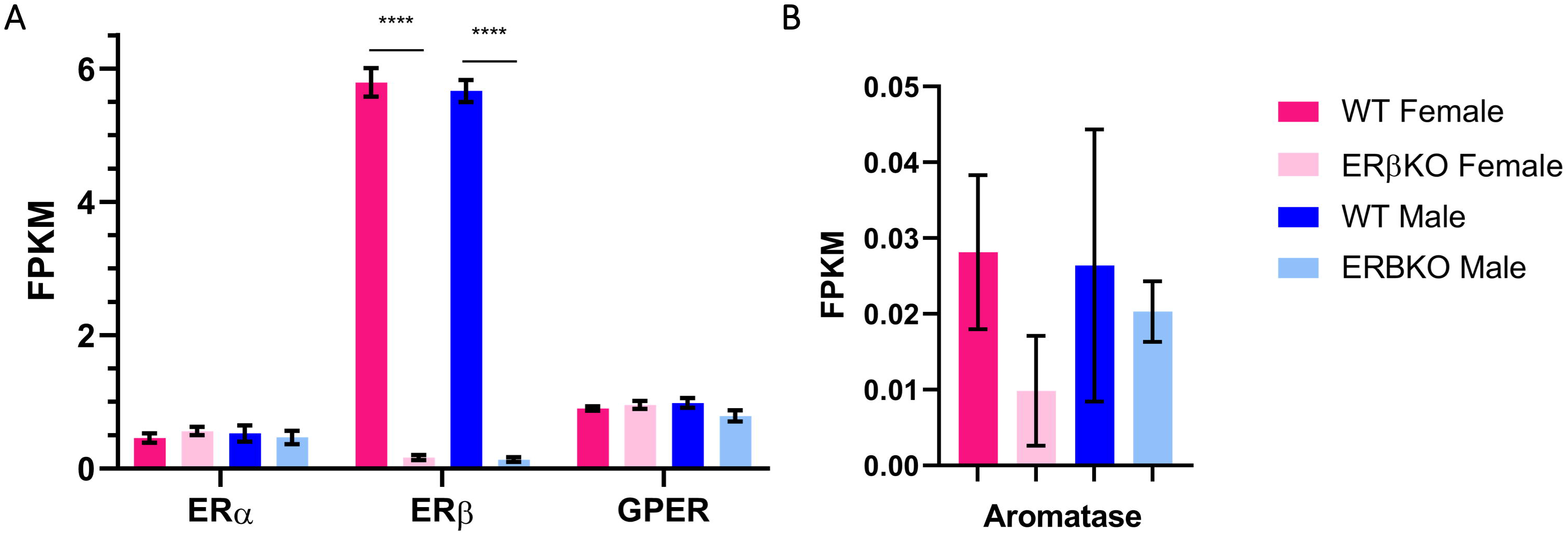
Expression levels of the three main estrogen receptors in the posterior cortex. **A.** FPKM values for the three estrogen receptor genes (*ERα, ERβ*, and *GPER*) averaged for each group (n = 4). A very low level of ERβ mRNA is likely present in the ERβKO mice because this line has a neo cassette introduced into exon 3, generating an early stop codon and no functional protein, but reads may map to exons 1 and 2 in the remaining mRNA fragment. ERβ expression in WT vs ERβKO animals was significantly different (p<0.001), but there were no significant differences in the expression of the other estrogen receptors. **B.** FPKM values for aromatase (*CYP19a1*) averaged for each group (n = 4). For both panels, significance was determined by the Wald test with Benjamini-Hochberg correction for multiple comparisons, and results are presented as mean ± SEM.

### Neither ERβ nor sex contribute to whole-transcriptome changes in the adult mouse cortex

To analyze global changes in the cortical transcriptome due to sex and ERβKO, we first performed principal component analysis on the normalized count data. Samples were separated along the first principal component by sex, but were not separable by genotype. Furthermore, 95% confidence interval probability ellipses plotted for each group along the first two principal components largely overlapped for ERβKO and WT, indicating that these groups are not distinct (Fig 3A). The correlation between samples, computed with Pearson’s correlation coefficient, was low, ranging between 0.34 and −0.43 excluding self-correlations, and was not higher within groups than between them (Fig 3B). Therefore, gene expression profiles cannot distinguish between groups. A heatmap of Z-scores for each gene showed no obvious distinctions between groups (Fig 3D). Unsupervised hierarchical clustering of samples, using both Pearson’s correlation coefficients (Fig 3C) and Z-scores (Fig 3E), showed that samples could not be clustered into groups based on sex or genotype. Importantly, estrous stage did not influence where individual females fell in either principal component or clustering. Overall, these data solidify our conclusions that male and female cortices do not have widely different patterns in gene expression, and that ERβKO does not have an appreciable effect on transcription in the cortex.

**Figure 3.**
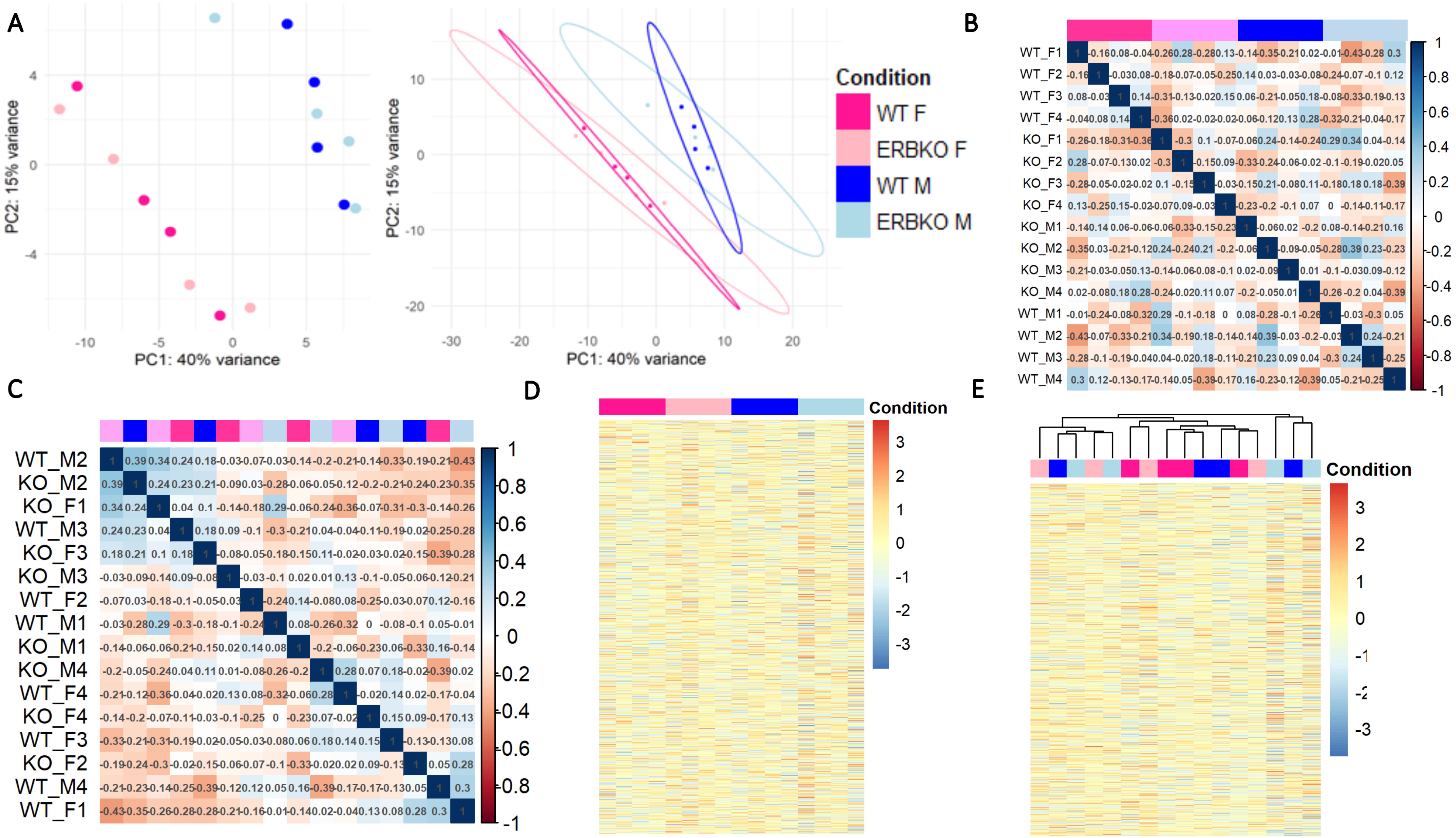
Analysis of whole-transcriptome changes in the posterior cortex. **A.** Principal component analysis of variance-stabilized count data for all samples. Sex is separated along the x-axis by PC1, while PC2 fails to separate samples by genotype along the y-axis. Ellipses show 95% confidence intervals calculated based on the probability distribution for each group. Color-coding of groups applies to all panels. **B.** Correlogram of Pearson’s correlation coefficient calculated for each sample against all other samples, range −0.43 – 0.34 excluding self-correlations. **C.** Unsupervised agglomerative hierarchical clustering of correlation coefficients into four clusters. **D.** Heatmap of Z-scores for the 15,546 genes with non-zero expression in all samples. Each column corresponds to one sample, and rows correspond to genes. **E.** Hierarchical clustering of Z-scores.

### There are few significant DEGs in male and female cortices

Although sex and ERβ did not contribute to a global regulation of gene expression in the cortex, this did not exclude the possibility that they may impact the transcription of a smaller number of genes, which may have been lost in the large-scale analyses. To address this possibility, we examined differential expression of genes identified in our dataset. We first looked at sex differences by comparing WT and ERβKO males and females. Histograms of the obtained adjusted p-values and log_2_ fold changes (Fig 4A and B) show that the overwhelming majority of p-values clustered around 1, and fold changes around 0, indicating very low differential expression between groups. However, we did find a small number of DEGs (adjusted p-value < 0.05). Six DEGs, all located on the X/Y chromosomes, were identified in the ERβKO female vs male comparison. These six DEGs plus 21 additional DEGs (27 total) were identified in WT females vs males (Fig 4C and D). Of the 21 genes with a sex difference only in WT animals, 17 were autosomal genes that displayed very small but statistically significant differences (Supplementary Data). A recent study by Vied et. al. reported over 900 sexually dimorphic DEGs in the WT C57BL/6J hippocampus (Vied et al., 2016). We compared our dataset from the WT female vs male comparison to the dataset generated by Vied et. al. in hippocampus, and although we found nearly 30,000 genes in common, including all the DEGs identified by Vied et. al., we identified only 9 common DEGs, all sex chromosome-linked (Fig 5).

**Figure 4.**
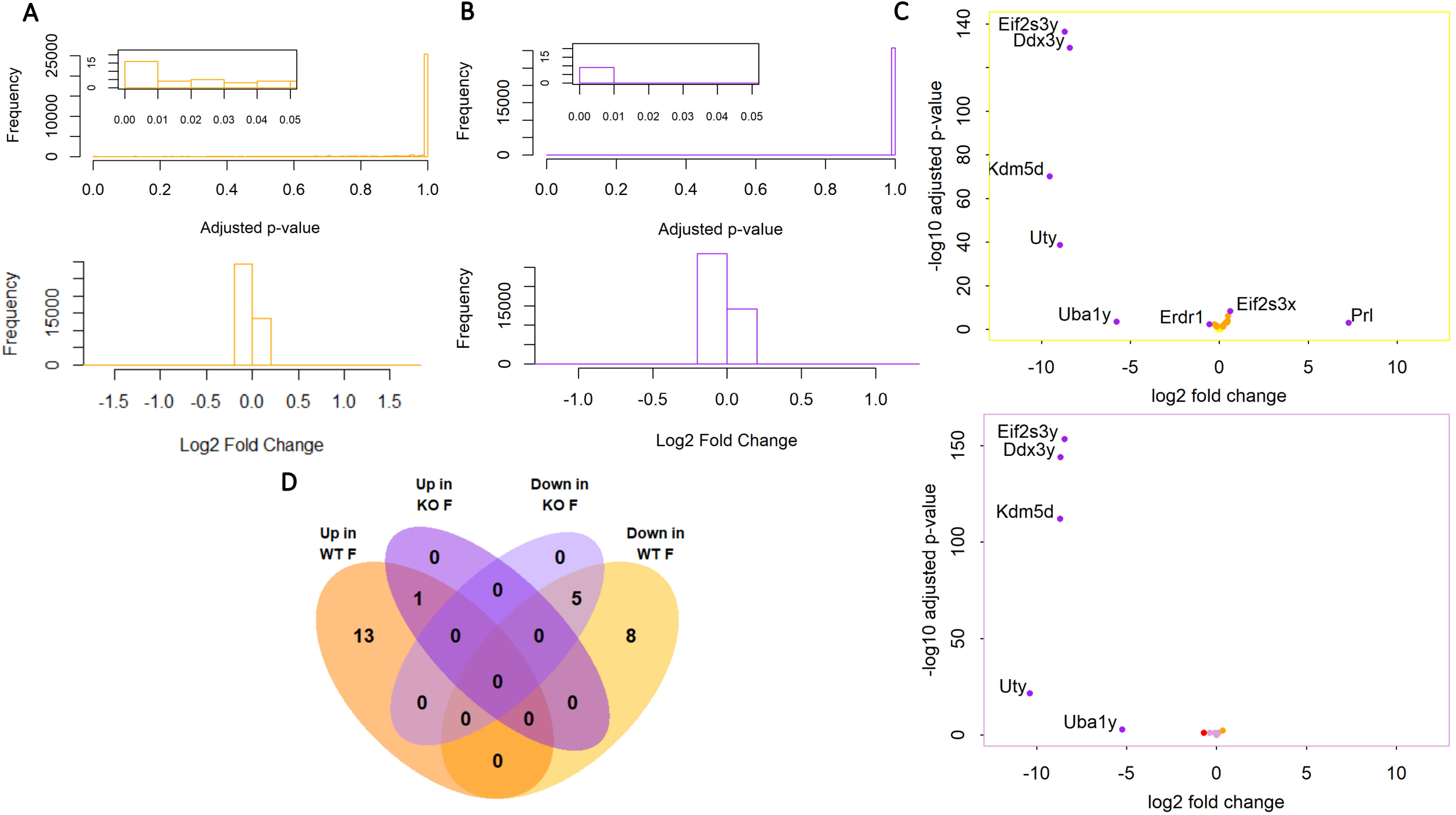
DEGs in males vs females. **A-B.** Histograms of adjusted p-values and log_2_ fold changes for the WT female vs male (shown in yellow, **A**) and ERβKO female vs male (shown in purple, **B**) comparisons, calculated by DESeq2. Insets show statistically significant p-values. **C.** Volcano plots showing −log_10_-transformed p-values plotted against log_2_ fold changes for each gene in the WT female vs male and ERβKO female vs male comparisons. Yellow points are significant (adjusted p-value < 0.05), red points have a log_2_ fold change greater than 50%, and purple points are significant and have a log_2_ fold change greater than 50%. **D.** Venn diagram showing the overlap between DEGs identified in the WT female vs male and ERβKO female vs male comparisons.

**Figure 5.**
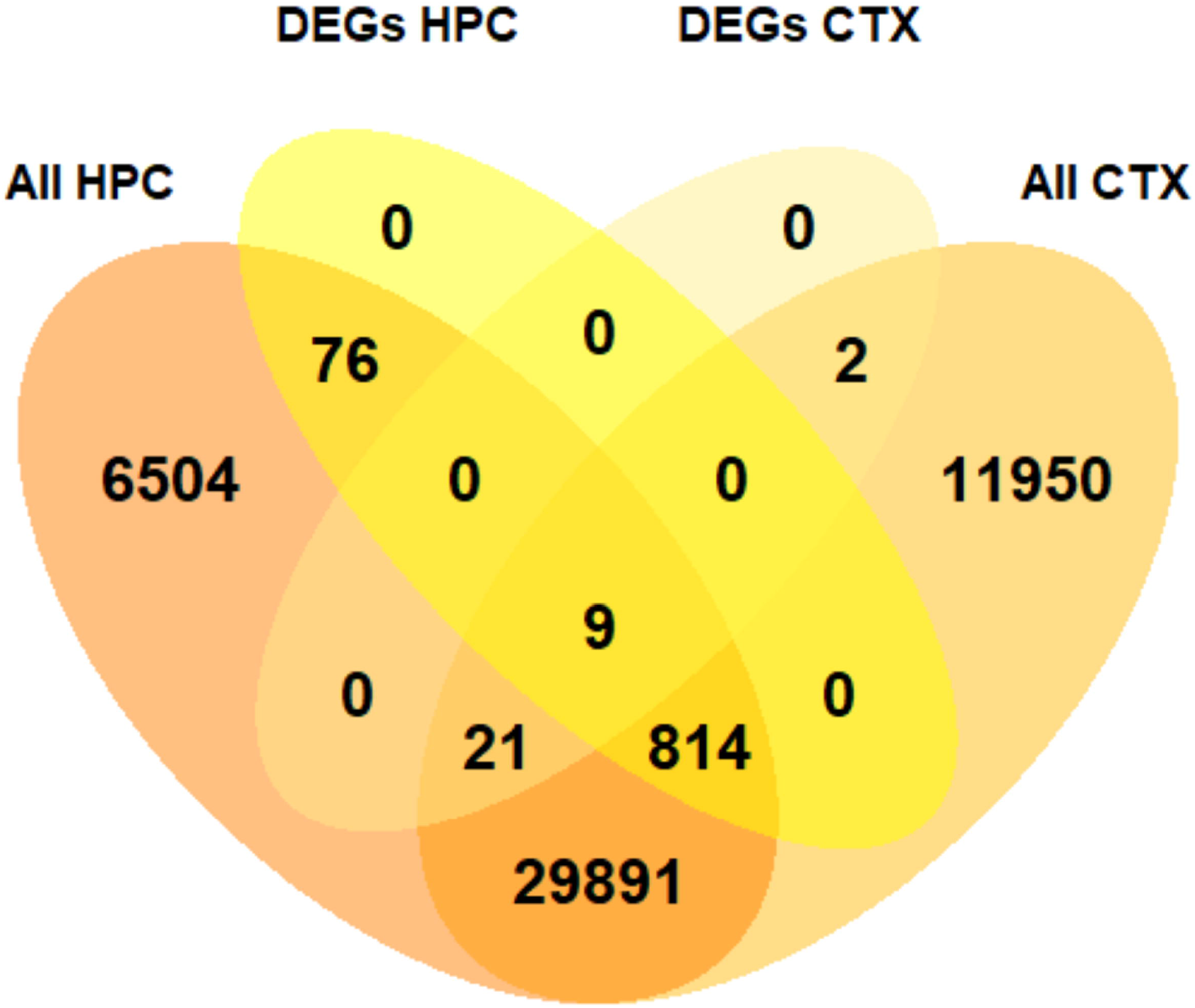
Comparison of genes with a sex difference in expression identified here in the posterior cortex and by Vied et. al. in the hippocampus. Venn diagram showing overlap between the DEGs in WT females vs males identified in this study in the cortex and those identified by Vied et. al. in the hippocampus (Vied et al., 2016).

### ERβ contributes to sex differences in the expression of cation channel genes

To elucidate the effects of ERβ on sex differences in cortical gene expression, we further investigated the genes with a significant sex difference in WT, but not ERβKO. As expected, a heatmap of the Z-scores of these genes (Fig 6A) shows that hierarchical clustering separates WT males and females completely, based on the expression of 20 out of the 21 genes that showed sex differences. Only 20 genes were used, because one gene had an expression level of 0 in two of the samples, so computing a Z-score not possible. We noticed that clustering of these genes produced two distinct subsets, regulated in a similar manner across groups. Cluster 1 contained 8 genes and its expression was lowest in WT females and highest in WT males, while cluster 2 contained 12 genes and was highest in WT females and lowest in males. Gene Ontology Enrichment revealed that cluster 2 was significantly enriched for genes with molecular functions related to cation channel activity (Fig 6B). Expression levels of the four genes from cluster 2 annotated with this function are shown in Fig 6C. Sex differences in the expression of three of these genes, *Lrrc55, Kcnq5*, and *Scn1a*, in WT but not ERβKO mice, were confirmed by qPCR (Fig 6D). *Slc24a2* was not analyzed by qPCR because of technical difficulties in amplification due to the existence of multiple isoforms of this gene.

**Figure 6.**
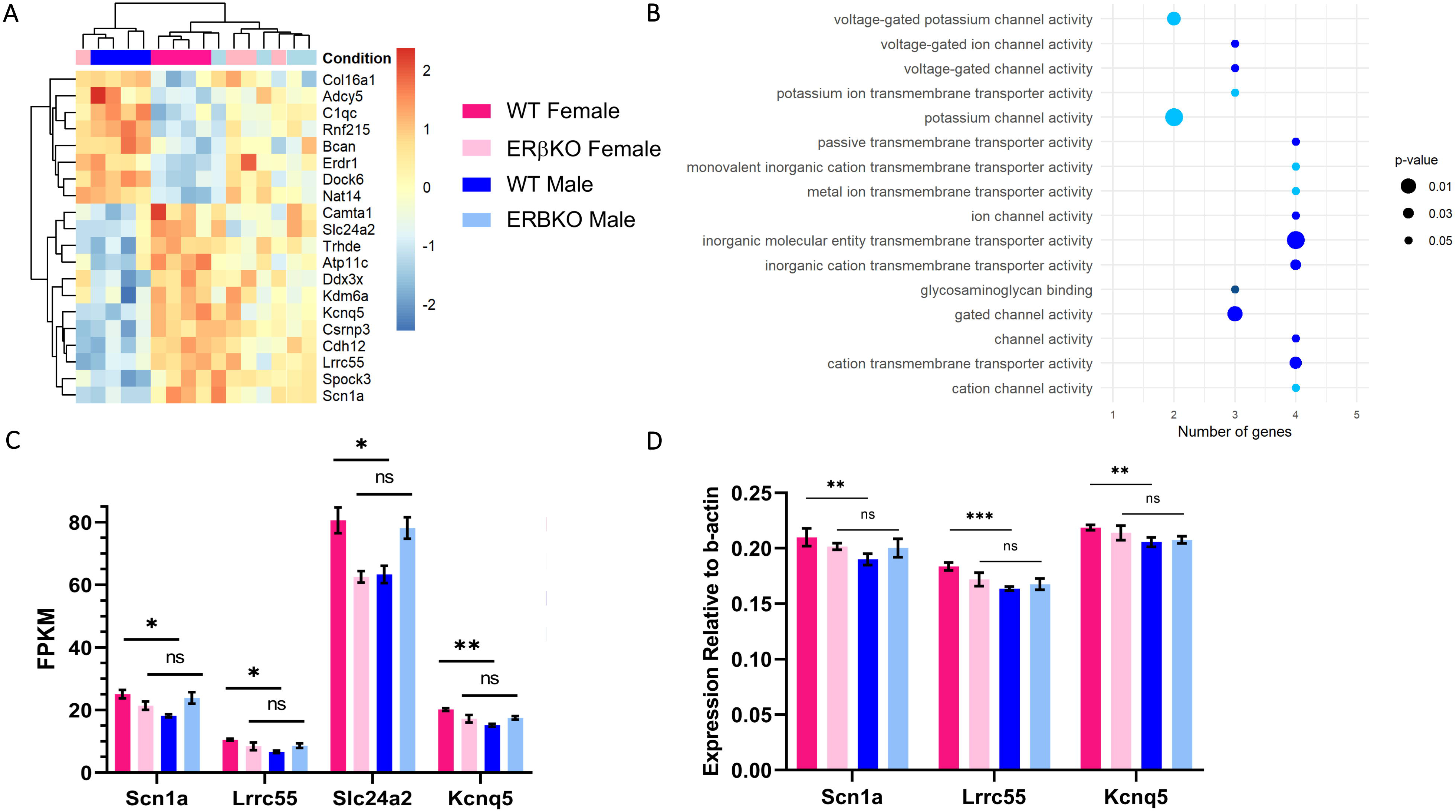
Gene ontology analysis of DEGs with a sex difference in WT but not ERβKO mice. **A.** Heatmap showing Z-scores for 20 of the 21 genes with a significant sex difference in WT but not ERβKO mice. The color-coded legend shown here in panel A applies to panels A, C, and D. **B.** List of significantly enriched Gene Ontology: Molecular Function categories for genes from Cluster 2, with number of genes in each category indicated on the x-axis. The size of the points corresponds to the p-value, and points of the same color belong to the same family of Gene Ontology terms. **C.** FPKM values for the four genes annotated with cation channel function averaged for each group (n = 4). All comparisons are significant in the Wald test with Benjamini-Hochberg correction for multiple comparisons for WT females vs males, p < 0.001 for Kcnq5 and p < 0.05 for all others, and not significant in ERβKO females vs males. Results are presented as mean ± SEM. **D.** Expression of *Scn1a, Kcnq5*, and *Lrrc55* by qPCR normalized to β-actin (*ACTB*) using the ∆Ct method, averaged for each group (n = 3). Significance was determined using t-tests with the Bonferroni method for correction for multiple comparisons. Results are presented as mean ± SEM.

### Meta-analysis of single-cell sequencing data indicates that the sexually dimorphic genes in the posterior cortex are mainly neuronal

While the only way to accurately quantify cell-type specific gene expression is with single-cell transcriptomics, this method is costly and technically challenging. To circumvent this issue, recent advances in computational methods have provided strategies for deconvolution of bulk RNA-sequencing datasets. One of these methods, multi-subject single cell expression reference (MuSiC), allows the experimenter to compare bulk sequencing data to a single-cell transcriptomics dataset obtained from the same tissue for an estimation of the proportions of cell types that contributed to the bulk RNA pool (X. Wang et al., 2019). We applied this method to our dataset and used single-cell sequencing of the C57BL/6J primary visual cortex as a reference (Tasic et al., 2016). We found that the majority (75%) of the RNA in our experiment most likely came from neurons (Fig 7A). Furthermore, we found that in the single-cell sequencing data from Tasic et. al. (Tasic et al., 2016), the four DEGs we identified annotated with the molecular function ‘cation channel activity’ were almost exclusively expressed in excitatory neurons (Fig 7B).

**Figure 7.**
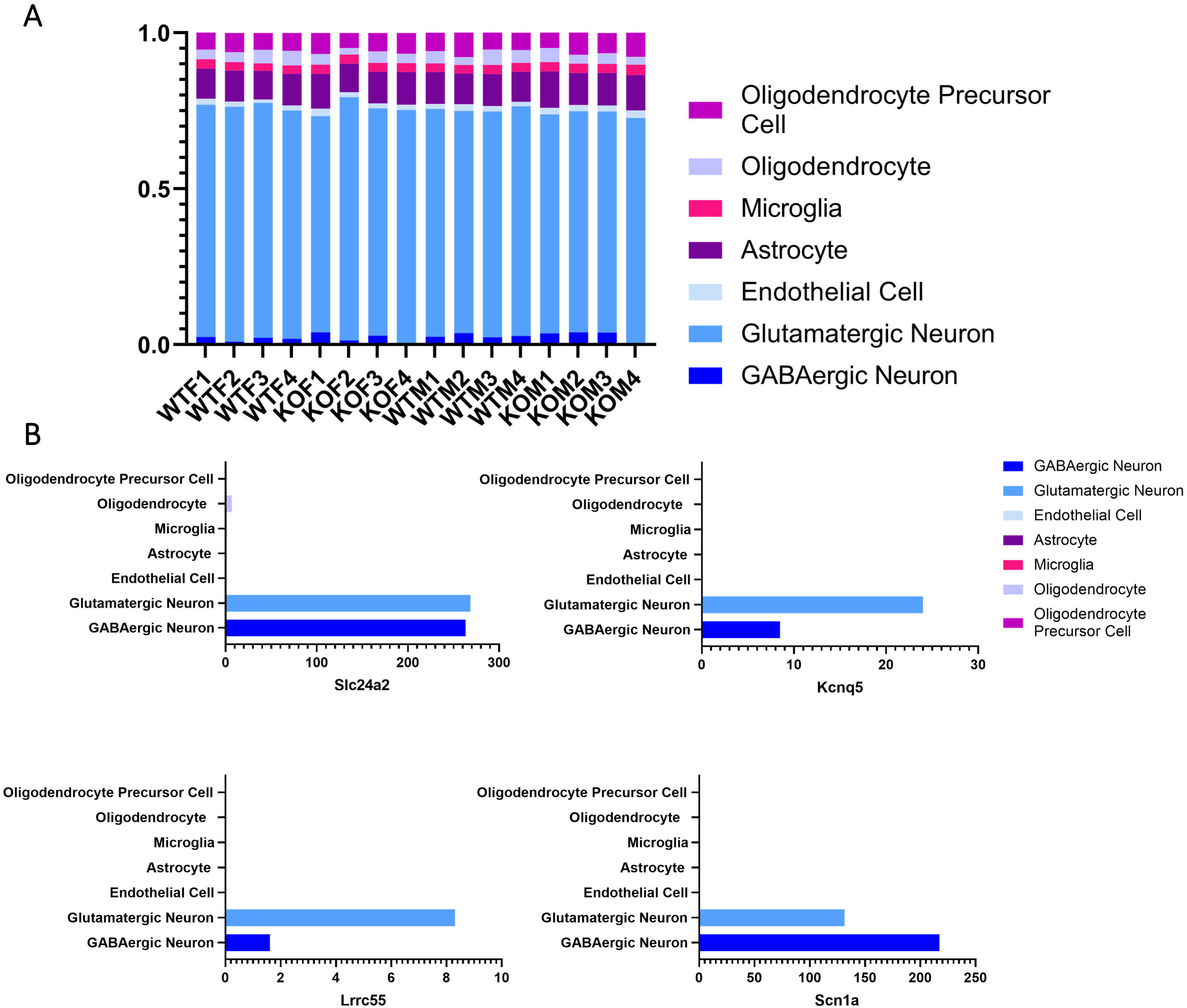
Estimation of cell type contributions to the RNA pool. **A.** Bar graph showing estimated proportional contribution of each cell type to our bulk RNA pool. **B.** Cell-type specific expression of the four DEGs with a sex difference in WT only, annotated with the molecular function ‘cation channel activity’. Expression levels calculated from normalized counts averaged across cell types, raw data and cell type annotation from Tasic et. al. (Tasic et al., 2016).

### Very few genes are differentially expressed in ERβKO cortices compared to WT in either sex

We next compared ERβKO to WT animals in each sex. Histograms of the p-values and fold changes show very few statistically significant DEGs (Fig 8A and B). Four significant DEGs were identified in both males and females, while four were uniquely found in males and one in females (adjusted p-value < 0.05) (Fig 8C) (Supplementary Data). Interestingly, the DEG unique to females, *Erdr1*, is the only gene that also shows a sex difference in WT, but not ERβKO. More genes were up-regulated, rather than down-regulated, in ERβKO animals (Fig 8D), suggesting that the transcriptional function of ERβ in the mouse cortex is limited to repressing the transcription of a small number of genes.

**Figure 8.**
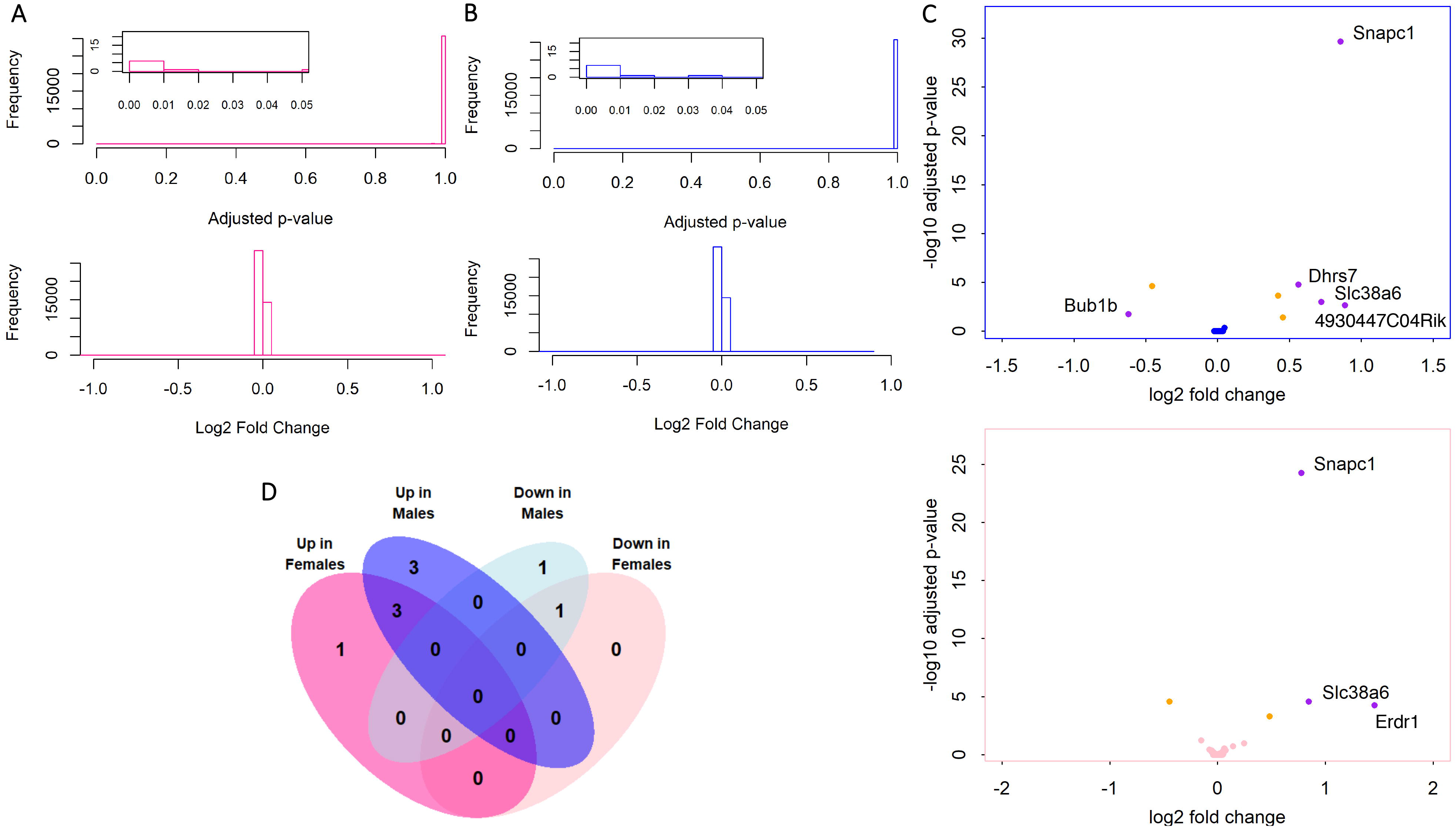
DEGs in WT vs ERβKO. **A-B.** Histograms of adjusted p-values and log_2_ fold changes for the female (shown in pink, **A**) and male (shown in blue, **B**) ERβKO vs WT comparisons, calculated by DESeq2. Insets show statistically significant p-values. **C.** Volcano plots showing −log_10_-transformed p-values plotted against log_2_ fold changes for each gene in the female and male ERβKO vs WT comparisons. Yellow points are significant (adjusted p-value < 0.05), red points have a log_2_ fold change greater than 50%, and purple points are significant and have a log_2_ fold change greater than 50%. **D.** Venn diagram showing the overlap between DEGs identified in the female and male ERβKO vs WT comparisons.

## Discussion

In the present study, we report sex- and ERβ-dependent gene expression profiles in the adult mouse posterior cortex. Surprisingly, we found very modest differences between sexes and genotypes. Principal component analysis could distinguish samples by sex, but not by genotype, and neither males and females nor ERβKO and WT animals could be clustered into groups based on their gene expression patterns. Loss of ERβ produced a very small number of statistically significant DEGs, 8 in males and 5 in females, of which 9 were uniquely identified genes and 4 were overlapping genes. We found 27 DEGs with a significant sex difference in WT mice, and interestingly the majority (21/27 genes) of these sex differences were lost in ERβKO animals. This suggests that ERβ regulates transcription of these genes in WT females, thus contributing to a sex difference in the expression of this limited number of genes in the posterior cortex.

The vast majority (176/181) of Y chromosome predicted protein coding genes were not classified as statistically significant DEGs, because they were lowly expressed in males and undetectable in females, and therefore failed to pass DESeq2’s low counts filter. Nevertheless, we identified five Y chromosome and one X chromosome DEGs in the ERβKO female vs male comparison. We found that these six genes plus three additional X chromosome genes, one gene expressed from both the X and Y chromosomes, and 17 autosomal genes were differentially expressed in WT females vs males. Interestingly, the DEG with a sex difference in WT animals located on the pseudoautosomal regions of both the X and Y chromosomes, *Erdr1*, is the only gene uniquely regulated in ERβKO vs WT females. *Erdr1*, a gene with unknown function in the brain, has also been shown to have a sex difference in expression in the cortex of neonatal mice (Armoskus et al., 2014).

We further analyzed the 21 DEGs with significant sex differences in the WT cortex that were lost in the ERβKO cortex. 17 of these are autosomal, suggesting that the sex difference in expression cannot be solely based on differential X/Y chromosome dosage in males and females. We therefore propose that ERβ may contribute to a sex difference in expression of these genes. We report that a subset of these genes with high expression in WT females is significantly enriched for genes annotated with molecular functions relating to cation channel activity. Studies have shown effects of sex on spontaneous neuronal firing rates in some brain regions (Blume et al., 2017; Dalpian, Rasia-Filho, & Calcagnotto, 2019; Osada, Nishihara, & Kimura, 1991), although the direction of these effects is highly dependent on brain region. Based on our results, we suggest that ion homeostasis may be a promising area for future research on functional sex differences in the brain. Furthermore, the effect of ERβ on neuronal excitability is largely unknown, and it will be interesting to determine whether ERβ contributes to sex differences in ion homeostasis and synaptic activity in the cortex.

The enrichment of cation channel DEGs led us to hypothesize that most genes with a sex difference in expression driven by ERβ are neuronal. The multi-subject single cell expression reference (MuSiC) algorithm allowed us to compare our bulk sequencing data to single-cell sequencing of the C57BL/6J primary visual cortex as a reference (Tasic et al., 2016) and estimate that 75% of the RNA in our experiment most likely came from neurons, consistent with estimations of the cell type contributions to bulk RNA pools from the brain reported by Tasic et. al. Furthermore, we examined the expression of the four DEGs with a sex difference in WT, but not ERβKO mice, annotated with the molecular function ‘cation channel activity’, in the single-cell sequencing data from Tasic et. al. These four DEGs are almost exclusively expressed in neurons, and ERβ is primarily detected in neurons as well (Tasic et al., 2016). This estimate suggests that the few sex differences in gene expression in the cortex mainly impact neurons.

While our study is limited to C57BL/6J mice, as this is the genetic background of the ERβKO line, it is interesting to note that this strain has been shown to have sexual dimorphism in anxiety and social behaviors (An et al., 2011). Additionally, the comparison of our findings in the posterior cortex with those reported by Vied et. al in the hippocampus (Vied et al., 2016) show that sex differences in the C57Bl/6J strain may be highly region-dependent. Interestingly, the 9 sex chromosome-linked DEGs found in common by us and Vied et. al. were also identified by this group as differentially expressed in the hippocampus in the 5 other strains examined (Vied et al., 2016). Several of these common DEGs located on the X chromosome are known to escape X-inactivation (Berletch, Yang, Xu, Carrel, & Disteche, 2011). This indicates that these genes may be part of a group of sexually dimorphic genes conserved across strains and brain regions. On the other hand, several studies have shown that sex differences in autosomal gene expression are highly tissue-specific (Norheim et al., 2019; Yang et al., 2006), and can be region-specific within one tissue (Nishida et al., 2005; Reinius et al., 2010). Thus, our results further highlight the importance of considering the effects of sex on transcription in a brain region-specific manner.

We chose to perform our study in randomly cycling females, in order to not limit the interpretation of our results to one specific estrous stage. In rodents, estrogen levels are low in metestrus, rise in diestrus, spike in proestrus, and fall back to low levels in estrus (Goldman, Murr, & Cooper, 2007). Since we found no females in proestrus at the time of sacrifice, estrogen levels were likely low and fairly consistent among our samples. However, we cannot rule out the possibility that sex differences in gene expression may exist in proestrus. Estrogen in the brain can originate from the gonads and cross the blood-brain barrier, and can also be synthesized by brain aromatase in both sexes (Ratner et al., 2019). Locally synthesized estrogen is detectable in the brain after ovariectomy, and can modulate cognition and behavior, even in the absence of circulating estrogen (Cornil, 2018; Vierk et al., 2012; Zhou et al., 2010). However, the relative contributions of circulating and locally synthesized estrogen to total levels in the brain remain unclear. In our dataset, aromatase (*CYP19a1*) expression was extremely low, and levels were not significantly different between any of the groups examined, suggesting that cortex-derived estrogen levels may have been similar across all samples in our experiment.

An interesting conclusion from our analysis is that loss of ERβ has minimal effects on gene expression in the posterior cortex of both sexes. While the canonical view of ERβ as mainly a hormone-responsive transcription factor has been questioned in recent years, many studies still focus on transcriptional regulation as the main mechanism of action for ERβ. In some tissues, including the female aorta (O’Lone et al., 2007) and the male motor cortex (Varshney et al., 2020), ERβ knockout has been shown to have significant effects on transcription. However, an ERβ mutant mouse in which only exon 3 is deleted, preventing the mutant protein from binding DNA, but still allowing it to participate in indirect transcriptional regulation and signaling, lacks some of the non-reproductive abnormalities seen in the ERβKO mouse. Although this study did not examine the brain, it suggests that direct ERE-binding may not be the primary mechanism of ERβ’s effects in peripheral tissues (Maneix et al., 2015). Several studies have shown that ERβ can repress AP-1 mediated transcription (C. Zhao, Dahlman-Wright, et al., 2010; C. Zhao, Gao, et al., 2010), suggesting that ERβ may function primarily as an indirect transcriptional regulator through AP-1. Our results, however, do not support this interpretation in the posterior cortex, and instead point to a mainly non-transcriptional mechanism for ERβ’s actions. ERβ has known effects on intracellular signaling cascades that can impact cognition, including the ERK and PI3K/Akt pathways (Sellers, Raval, & Srivastava, 2015). Furthermore, several studies have shown rapid signaling effects of ERβ on cortical neuron spine formation and synaptogenesis (K. J. Sellers et al., 2015; Srivastava, Woolfrey, Liu, Brandon, & Penzes, 2010), and inhibiting the ERK pathway can block ERβ’s neuroprotective effects in hippocampal neurons (L. Zhao & Brinton, 2007). Our data align with this framework, supporting a non-transcriptional mechanism of action for ERβ in the posterior cortex.

The present study raises a number of questions. First, if functional sex differences in the posterior cortex are not due to transcriptional differences, what are they caused by? Investigating sex differences in alternative splicing, posttranslational modifications, local protein synthesis, protein localization, and/or protein-protein interactions could provide answers. Moreover, if the effects of ERβ in the posterior cortex are not transcriptional, what are they due to? Evidence described above suggests that ERβ can act by regulating signaling pathways and protein-protein interactions. Investigating these non-transcriptional mechanisms in the cortex could provide insight into the region-specific effects of ERβ on sex differences in the brain.

## Acknowledgements

We would like to acknowledge the Weill Cornell Genomics Resources Facility, headed by Dr. Jenny Zhaoying Xiang. Funding for this work was provided by the NIH-NINDS R01NS095692.

